## Supplementary Material for "The Bardet-Biedl protein Bbs1 controls photoreceptor outer segment protein and lipid composition"

### Supplementary data

Suppl. Fig. 1: **Synteny of the zebrafish *bbs1* locus**

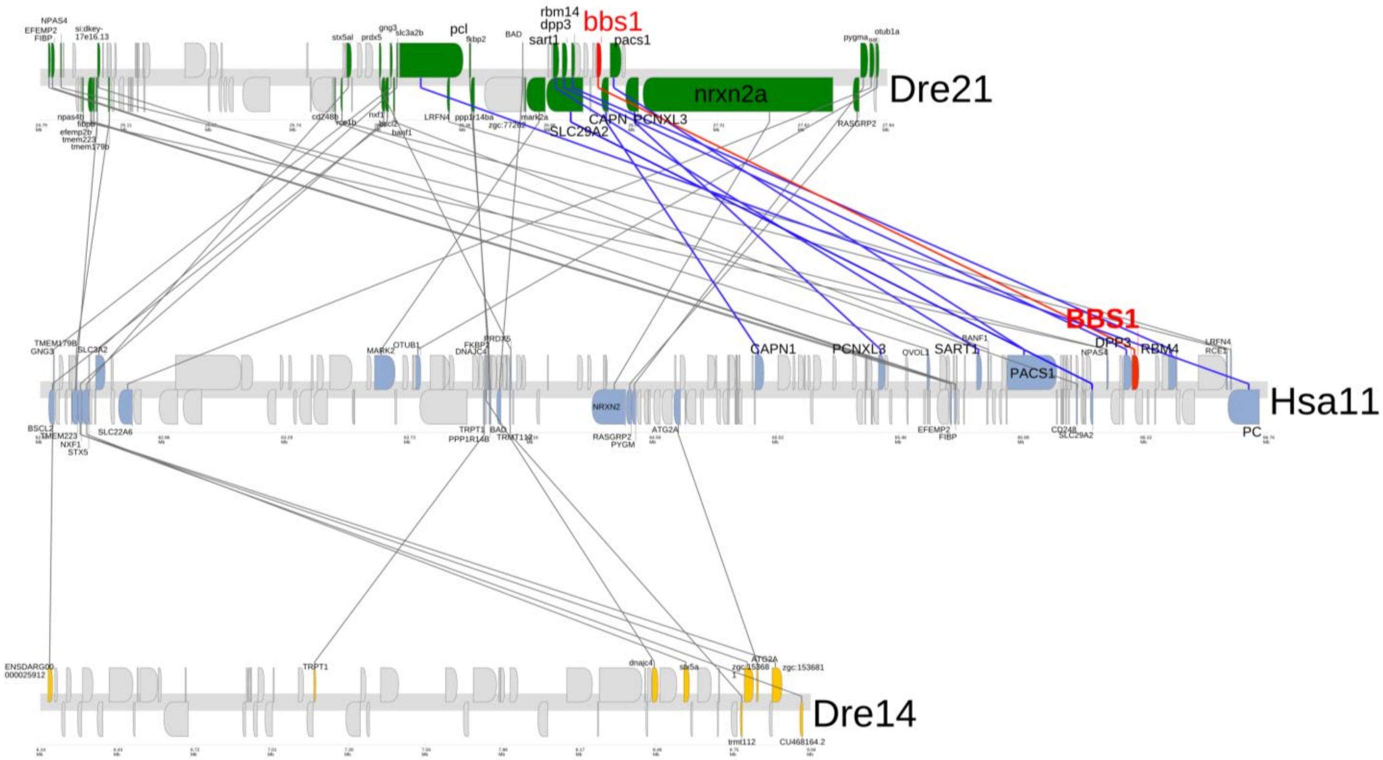


Genes flanking *BBS1* located on human chromosome 11 are found on zebrafish chromosomes 21 and 14. BBS1 genes are highlighted in red. The localization within the chromosome is given in the scale bars below the chromosomes. Orthologous genes between human (blue) and corresponding genes on zebrafish chromosomes 21 (green) and 14 (yellow), are depicted. The grey lines linking corresponding genes indicate the relative position of the genes on the chromosome and point out the single zebrafish orthologue. Orthologous genes directly flanking the human BBS1 locus are highlighted by dark blue lines. Note that most of these genes are found around the *bbs1* locus of zebrafish chromosome 21. *Dre* Danio rerio, *Hsa* Homo sapiens.

Suppl. Fig. 2: **Protein homology between human and zebrafish Bbs1**

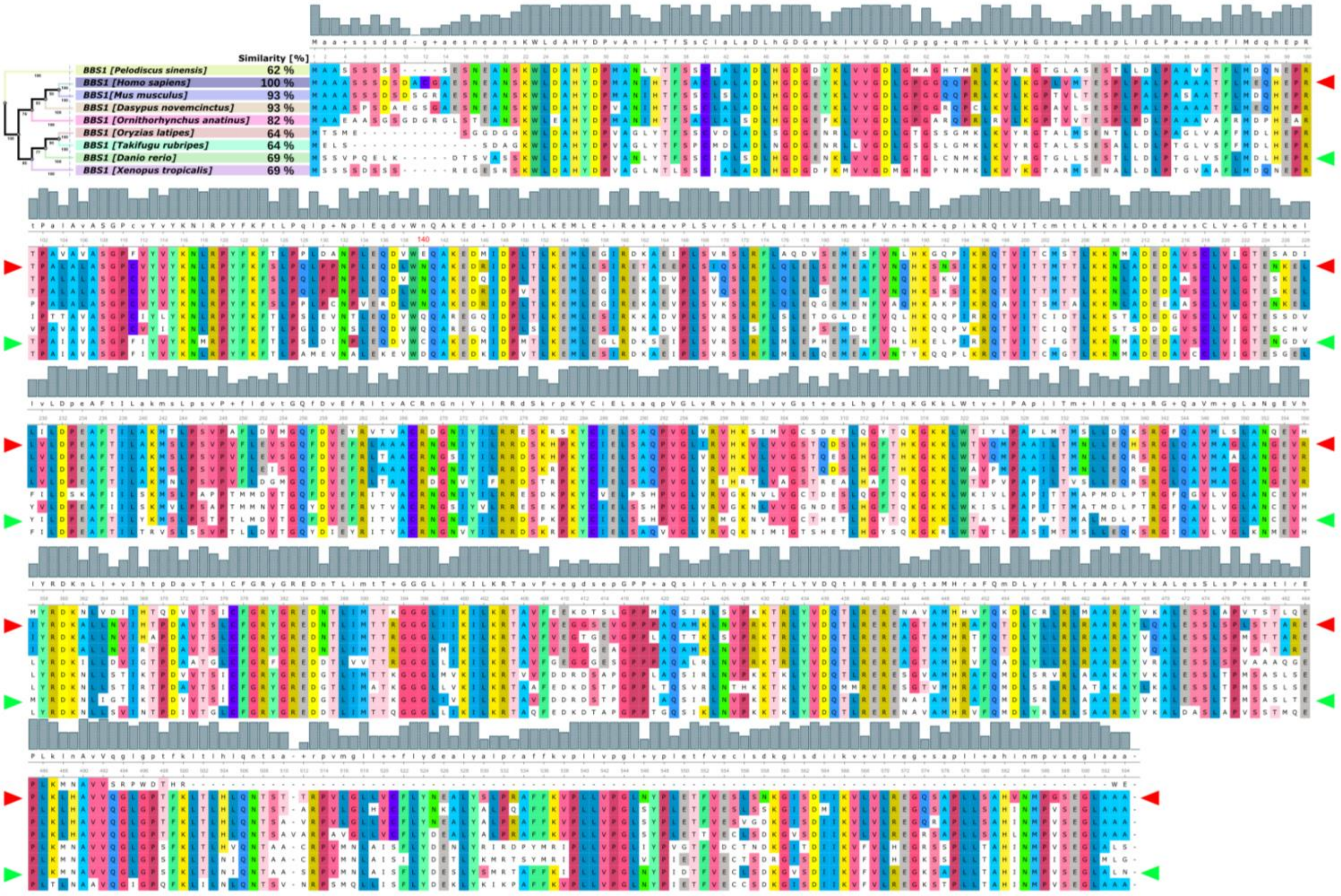


Amino acid sequences of the following species were aligned using Unipro UGENE (vers. 36.0), configured for high accuracy: Human *(Homo sapiens)*, mouse *(Mus musculus)*, zebrafish *(Danio rerio)*, torafugu *(Takifugu rubripes)*, medaka *(Oryzias latipes), armadillo (Dasypus novemcinctus), Chinese softshell turtle (Pelodiscus sinensis) platypus (Ornithorhynchus anatinus) and xenopus (Xenopus tropicalis).* Conservation is displayed as a bar graph (grey boxes) and the most conserved amino acid is the consensus (above sequence alignment at a given position). Conservation of the amino acid sequence between human and zebrafish Bbs1 is 69%. The red arrowhead points to the human sequence and the green arrowhead to the zebrafish sequence.

Suppl. Fig. 3: **Lack of opsin mislocalization despite progressive retinal dystrophy**

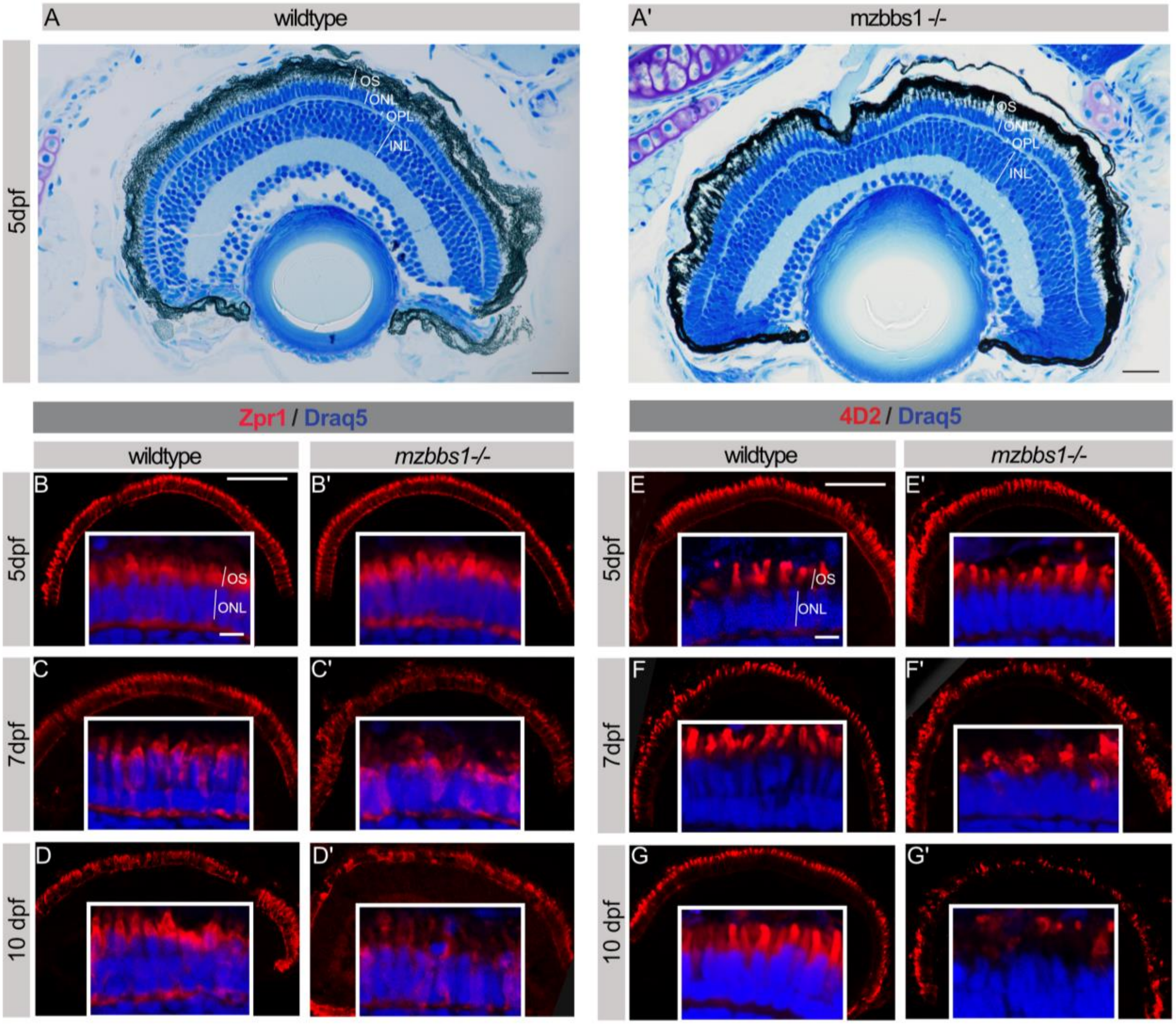


(**A, A'**) Semi-thin plastic section of *mzbbs1*$^{-/-}$ mutant (**A'**) and control eyes (**A**) stained with Richardson solution at 5 dpf are indistinguishable from each other. Retina lamination is unaffected by the mutation with respect to the overall morphology. The INL, ONL and OPL thickness in mutants are comparable to controls. (**B-D'**) Immunohistochemistry staining of red-green cones using zpr1 (red) reveals no morphological changes at 5 dpf (**B, B'**). Starting at 7 dpf (**C, C'**) slight morphological abnormalities are observed that progress over time and at 10 dpf (**D, D'**) the OS layer is substantially thinner and disorganized. (**E-G'**) Immunohistochemistry of opsins using the 4D2 antibody (red) shows no opsin mislocalization at 5 dpf (**E, E'**), 7 dpf (**F, F'**) or 10 dpf (**G, G'**) despite the severe morphological changes. Draq5 (blue) was used to counter stain the nuclei. Scale bars: (A, A'): 20 µm, (B-G'): 50 µm, inserts: 10 µm; Abbreviations: *OS*, outer segment; *ONL*, outer nuclear layer; *OPL*, outer plexiform layer; *INL*, inner nuclear layer.

Suppl. Fig. 4: **Increased photoreceptor cell death in *mzbbs1*$^{-/-}$ mutants at 10 dpf**

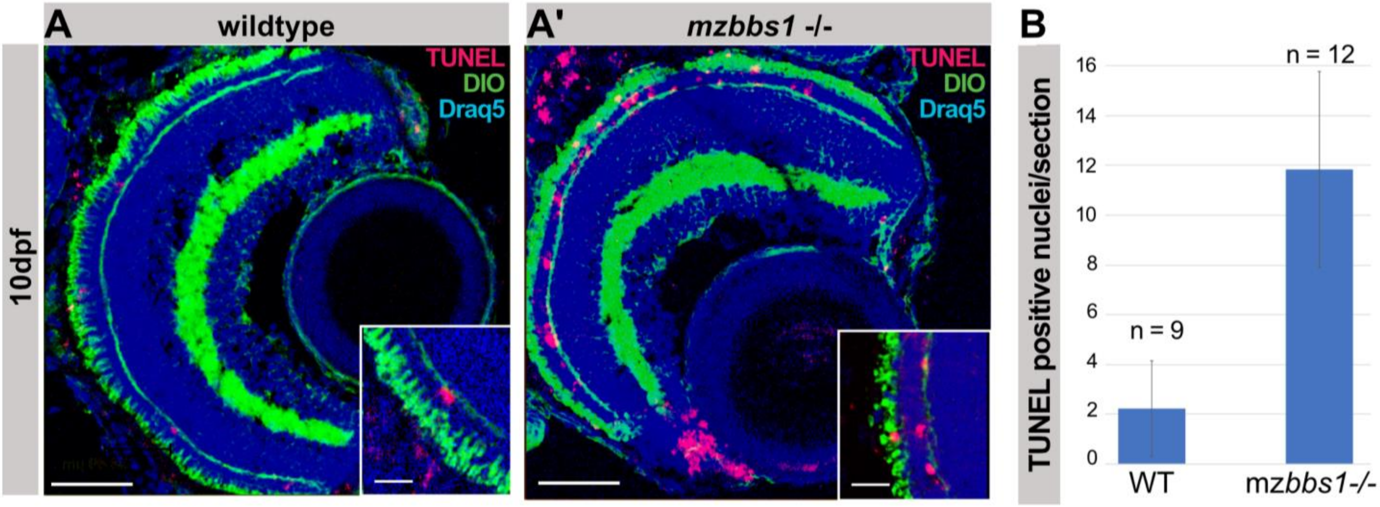


Apoptotic cells are labeled using the ApopTag® Red In Situ Apoptosis Detection Kit in control **(A)** and *mzbbs1* mutants **(A')**. Quantification of apoptotic cells per section is shown in **(B).** Error bars mark the standard deviation. Scale Bar: (**A, A'**) 47 µm, insert: 12 µm.

Suppl. Fig. 5: **Slowly progressive retinal dystrophy in zygotic *bbs1* mutants**

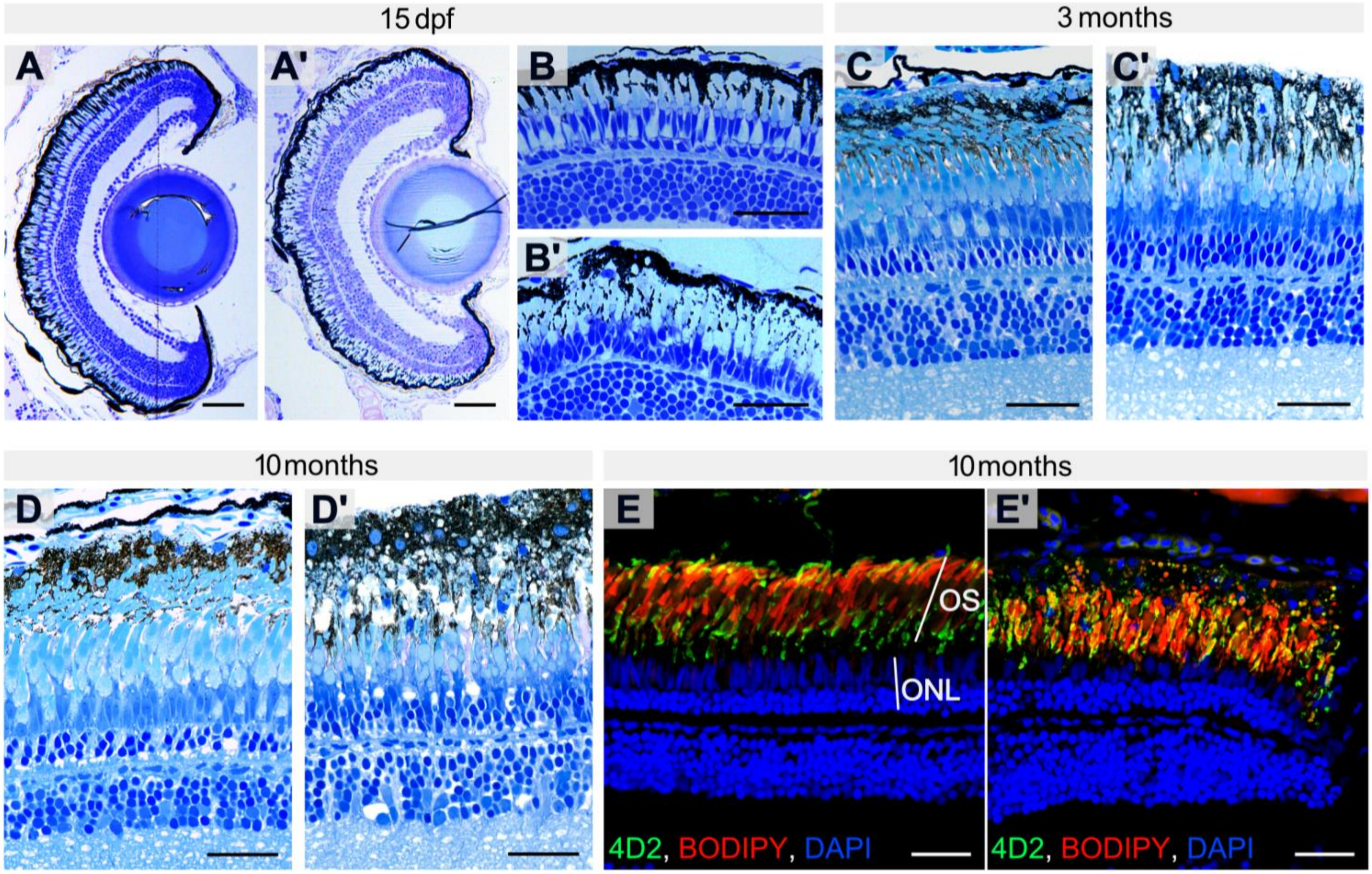


(**A-D'**) Semi-thin plastic sections of *zgbbs1*$^{-/-}$ mutants at 15 dpf (**A', B'**), 3 months (**C'**) and 10 months (**D'**) compared to controls (**A-D**) show a slow progression of retinal dystrophy. At 3 months post fertilization (mpf) some healthy-looking outer segments are still present in mutant retinas (**C'**). At 10 mpf the mutant OSs layer is severely disrupted (**D'**) and looks degenerated compared to control (**D**). Nuclei are observed within the mutant OS layer at 10 months, suggesting an invasion of microglia or abnormal RPE (**D'**). (**E, E'**) Immunostaining of opsins using 4D2 (green) reveals no opsin mislocalization in the IS or ONL in the retina of 10 mpf mutants (**E'**). The sections are co-stained with the membrane specific dye BODIPY to label OSs (red) and DAPI for nuclei (blue). Scale bar: (A, A'): 50 µm; (B-E'): 30 µm; Abbreviations: *OS* outer segment, *ONL* Outer nuclear layer

Suppl. Fig. 6: **Zygotic *bbs1* mutants show a slow decrease in visual function**

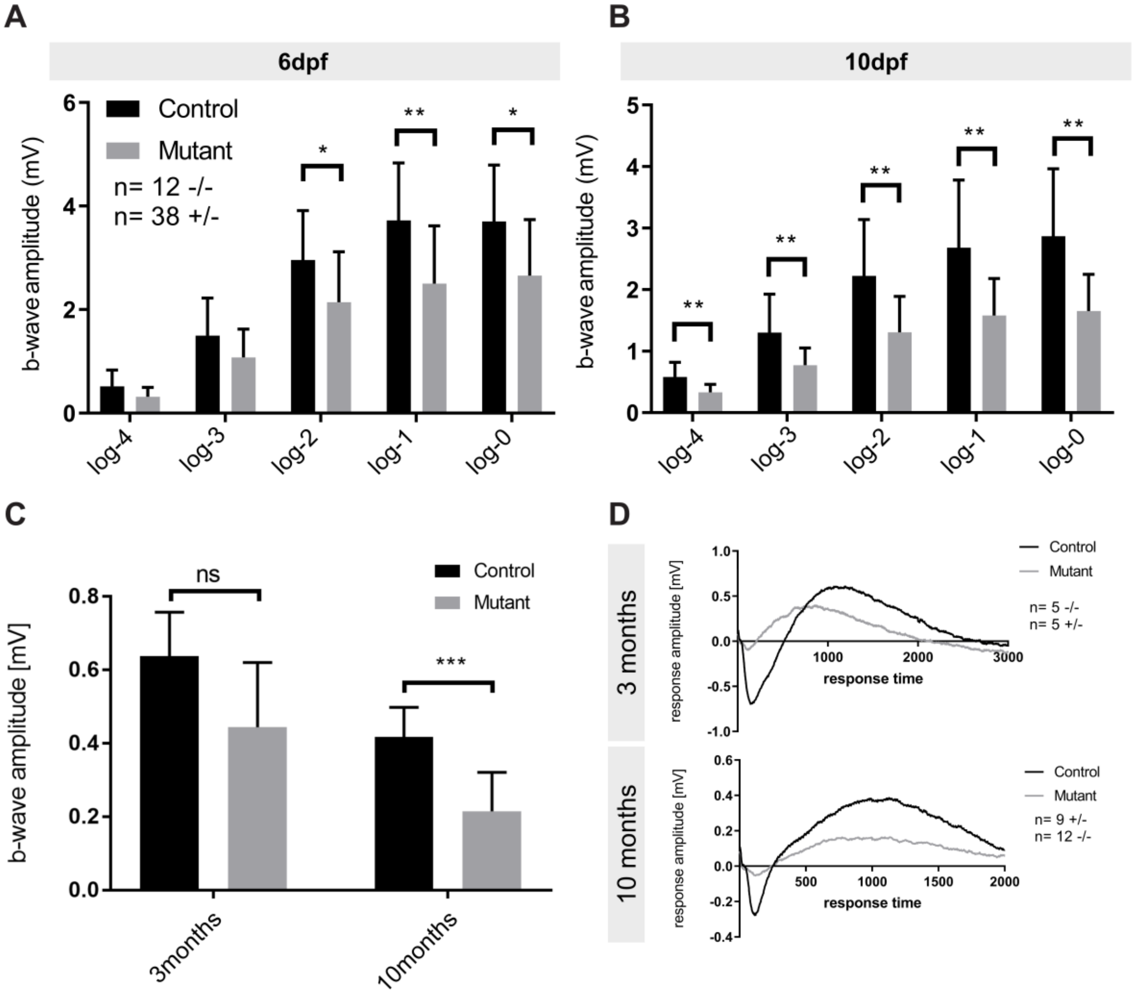


The functional phenotype of *zgbb1* mutants was assessed using ERG in larvae at 6 and 10 dpf as well as in adults at 3 and 10 months post fertilization. (**A-B**) Bar plots of the maximum b-wave amplitude by electroretinography (ERG) shows a significantly decreased response to light in *zgbbs1*$^{-/-}$ mutants for high light intensities (log-0 to log-2) at 6 dpf (**A**) and for all light intensities (log-0 to log-4) at 10 dpf (**B**). Unpaired two-tailed multiple T-test; Sig: ns= FDR(q-value)>0.05; *= FDR(q-value)<0.05; **= FDR(q-value)<0.01; Error bars show standard deviation. For more detailed statistics, please see **suppl. Table 5**. (**C**) Bar plot of the ERG response of adult fish at 3 and 10 months, at the highest light intensity (log-0 corresponds to 24’000µW/cm$^2$). Unpaired two-tailed multiple T-test, Holm-Sidàk adjusted; Sig: ns=adj.p-value=0.077, ***= adj.p-value <0.001, Sample size (n=5 WT, n=5 Mut eyes at 3 months & n=8 WT, n=11 Mut eyes at 10 months); Error bars show standard deviation. (**D**) Average ERG response curve for wt (black) and mutant (grey) at 3 months in top panel and 10 months in bottom panel. The lack of statistical significance at 3 months is likely explained by the technical difficulty of measuring ERGs in adult zebrafish, which limits the number of animals that can be measured. The clearly reduced average curve and the altered shape of the curve (shorter time to reach the maximum response) at this stage (**D**) support a visual deficit at 3 months already. Abbreviations: *ERG* Electroretinography.

Supp. Fig. 7: **Expression levels of BBSome component in the adult retina**

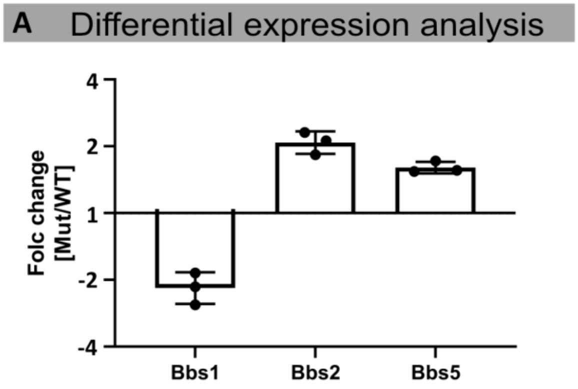


**B** Primer table

| Name | Primer | Sequence 5'-3' |
|---|---|---|
| BBS1_Frw | For. | CTC CTC ATG CAT TGC TCT TTC AG |
| BBS1_Rev | Rev. | CAC GAT ACA CCT TCA GCT TCA TG |
| BBS2_Frw | For. | AGA TGG CAG AAA ATG AGA CAG TCA C |
| BBS2_Rev | Rev. | TTT GAC TTG ATC CTC CAG TAG CG |
| BBS5_Frw | For. | CGG AGA TAG AGG GAG ACT TCT GG |
| BBS5_Rev | Rev. | GTA TCC CAC AGA CAG GTT GAC TC |
| g6pd_Frw | For. | CTG GAC CTG ACC TAC CAT AGC AG |
| g6pd_Rev | Rev. | AGG CTT CCC TCA ACT CAT CAC TG |

The expression levels of *bbs1*, *bbs2* and *bbs5* in adult retina of *zgbbs1*$^{-/-}$ mutants were assessed using real time PCR. (**A**) Expression levels were normalized to the expression of the housekeeping gene *g6pd* and fold changes were assessed between wildtype and mutants. Error bars show min/max values. Sample size: (n=3 Ctrl, n=3 Mut). Note the decrease in Bbs1 levels and the lack of downregulation of Bbs2 and Bbs5 transcripts. (**B**) Primer sequences used for the real time experiment.

Suppl. Fig. 8: **Proteomic analysis on isolated OSs**

**A** Mouse vs. Zebrafish proteome

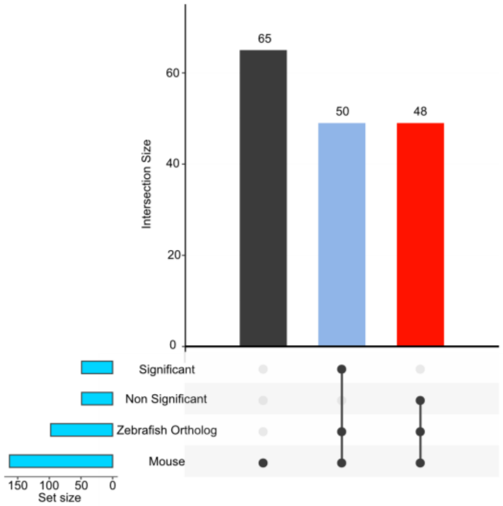


**B** Protein associated with lipid/lipid metaobolismus

| Associated with | Name | Description | UniprotID | FC | adj p.value |
|---|---|---|---|---|---|
| Cholesterol/ Retinoid transport | LOC565099 | NPC intracellular cholesterol transporter 2 | B3DFW4 | 3.3 | 1.69E-02 |
| | erlin1 | ER lipid raft associated 1 | Q58EG2 | 1.3 | 4.02E-02 |
| | scp2b | sterol carrier protein 2b | Q5XJS3 | 2.1 | 3.31E-04 |
| | erlin2 | cholesterol 25-hydroxylase like 3 | A3QK16 | -1.4 | 1.83E-04 |
| | abca4a | ATP-binding cassette, sub-family A (ABC1), member 4a | E9QD15 | -1.2 | 4.08E-02 |
| | abca4b | ATP-binding cassette, sub-family A (ABC1), member 4b | F1R045 | -1.3 | 2.06E-04 |
| LDL/HDL | apoeb | apolipoprotein Eb | O42364 | 2.4 | 5.78E-04 |
| | lrp1aa | low density lipoprotein receptor-related protein 1Aa | A0A0H2UKL4 | 2.5 | 3.39E-21 |
| | apoa4b.1 | apolipoprotein A-IV b, tandem duplicate 1 | F1QHR0 | 1.2 | 0.00E+00 |
| | lrpap1 | low density lipoprotein receptor-related protein associated protein 1 | A0A2R8QMD9 | 2.7 | 8.35E-12 |
| | apoc1 | apolipoprotein C-I | E9QFX0 | 1.9 | 9.00E-02 |
| | apoa1b | apolipoprotein A-Ib | A0A0R4IKF0 | 2.3 | 5.20E-04 |
| | apoa1 | apolipoprotein A-Ia | O42363 | 1.6 | 1.02E-02 |
| | scarb2a | scavenger receptor class B, member 2a, intreacellular lipid transport ; https://p | Q8JGR8 | 1.4 | 1.26E-01 |
| Lipid metabolism enzymes | ppt1 | palmitoyl-protein thioesterase 1 (ceroid-lipofuscinosis, neuronal 1, infantile) | F1R8J6 | 2.8 | 8.75E-12 |
| | pnpla8 | patatin-like phospholipase domain containing 8 | F1RE62 | -1.0 | 9.37E-01 |
| | pitpnaa | phosphatidylinositol transfer protein, alpha a | Q803U3 | 3.3 | 8.70E-12 |
| | inpp5ka | inositol polyphosphate-5-phosphatase Ka | A0A0R4IWP4 | 4.7 | 1.97E-20 |
| | pip4k2cb | phosphatidylinositol-5-phosphate 4-kinase, type II, gamma b | F1QRI7 | 3.2 | 1.78E-02 |
| | phospho2 | phosphatase, orphan 2 | A9JRB9 | 3.0 | 0.00E+00 |
| | chdh | choline dehydrogenase | E7EY13 | 2.1 | 1.17E-03 |
| | ppp2r2aa | protein phosphatase 2, regulatory subunit B, alpha a | X1WDS4 | 1.8 | 0.00E+00 |
| | pcyox1 | prenylcysteine oxidase 1 | A0A0R4I9Q1 | 1.8 | 3.82E-06 |
| | smpdl3a | sphingomyelin phosphodiesterase, acid-like 3A | F1Q7D3 | 1.5 | 0.00E+00 |
| | sacm1la | SAC1 like phosphatidylinositide phosphatase a | A4VCH0 | 1.5 | 5.61E-04 |
| | lmf2b | lipase maturation factor 2b | F1Q4Q4 | 1.4 | 0.00E+00 |
| | cpt2 | carnitine palmitoyltransferase 2 | B2GQ15 | 1.3 | 4.84E-02 |
| | cds1 | CDP-diacylglycerol synthase (phosphatidate cytidylyltransferase) 1 | A5PN44 | -1.7 | 0.00E+00 |
| | acox1 | acyl-CoA oxidase 1, palmitoyl | F1R3V9 | 3.2 | 2.13E-03 |
| | synj1 | synaptojanin 1 ; phosphatase | E9QHF3 | -1.3 | 1.47E-02 |
| Fatty acid | fabp11b | fatty acid binding protein 11b | Q503X5 | 5.7 | 6.88E-14 |
| | fabp1a | fatty acid binding protein 1a, liver | Q1AMT3 | 3.5 | 2.01E-04 |
| | fabp7a | fatty acid binding protein 7, brain, a | Q9I8N9 | 2.4 | 5.71E-10 |
| | fasn | Fatty acid synthetase | E7F5V3 | 1.2 | 3.96E-02 |
| | acsl2 | acyl-CoA synthetase long chain family member 2 | E7F351 | 1.5 | 1.34E-02 |
| Lipid binding | tollip | toll interacting protein | A0A0R4IZ33 | 1.8 | 6.86E-04 |
| | sec14l8 | SEC14-like lipid binding 8 | A2BIR0 | 1.5 | 7.68E-03 |
| | hsdl2 | hydroxysteroid dehydrogenase like 2 | Q6P5L8 | 1.5 | 8.72E-03 |
| | vtg1 | vitellogenin 1 | Q1LWN2 | 3.9 | 0.00E+00 |
| | vtg7 | vitellogenin 7 | Q1MTC6 | 6.5 | 0.00E+00 |
| PI/PL binding | vil1 | villin 1 | F1QVU3 | 2.7 | 6.53E-31 |
| | tulp1b | TUB like protein 1b | F8W3P9 | 2.7 | 1.11E-04 |
| | anxa2a | annexin A2a | Q6P603 | 3.8 | 3.06E-05 |
| | anxa3b | annexin A3b | Q5U369 | 4.8 | 1.57E-05 |
| | anxa4 | annexin A4 | Q804G7 | 1.8 | 8.79E-08 |
| | anxa5b | annexin A5b | Q6P0V8 | 2.8 | 0.00E+00 |
| | Plscr1 | si:ch73-206p6.1; Phospholipid scramblase 1 | X1WGC6 | 3.6 | 1.92E-11 |

(**A**) Graph illustrating the results from a direct comparison between the outer segment proteome of *Bbs17* mutant mouse (Datta et al 2015) and our *bbs1* zebrafish dataset. Out of the 146 proteins that were significantly enriched in the *Bbs17* mouse OSs, we found for 81 a zebrafish orthologue in our data set. Because the zebrafish genome has an additional genome duplication, in some cases we found several possible orthologues for a single mouse protein. Therefore, we found a total of 98 zebrafish proteins that are potential orthologues of the 81 mouse proteins. 50 proteins were significantly different (adj. pval <0.05) in both datasets and 48 zebrafish orthologues were only significant in the BBS17 mouse dataset but not in the bbs1 zebrafish data set. However, it is important to note, that out of the 16 mouse proteins where we found several zebrafish orthologues, in 8 cases the abundance of one paralogue was significantly altered in our data set and the other paralogue was not, therefore the second, not significantly altered paralogues inflates the total number of "non-significant" proteins in this comparison. For 65 mouse proteins, we were not able to find an orthologue in our data set. The comparison demonstrates a substantial overlap between our dataset and the dataset from *Bbs17* mouse (**see Supp. Table 3**). (**B**) Table describing the large number of lipid associated proteins that were found to be enriched in the zebrafish *bbs1* mutant OSs. We listed all the proteins found and grouped them according to their properties or function.

Suppl. Fig. 9: **Over-representation analysis of the proteomics dataset**

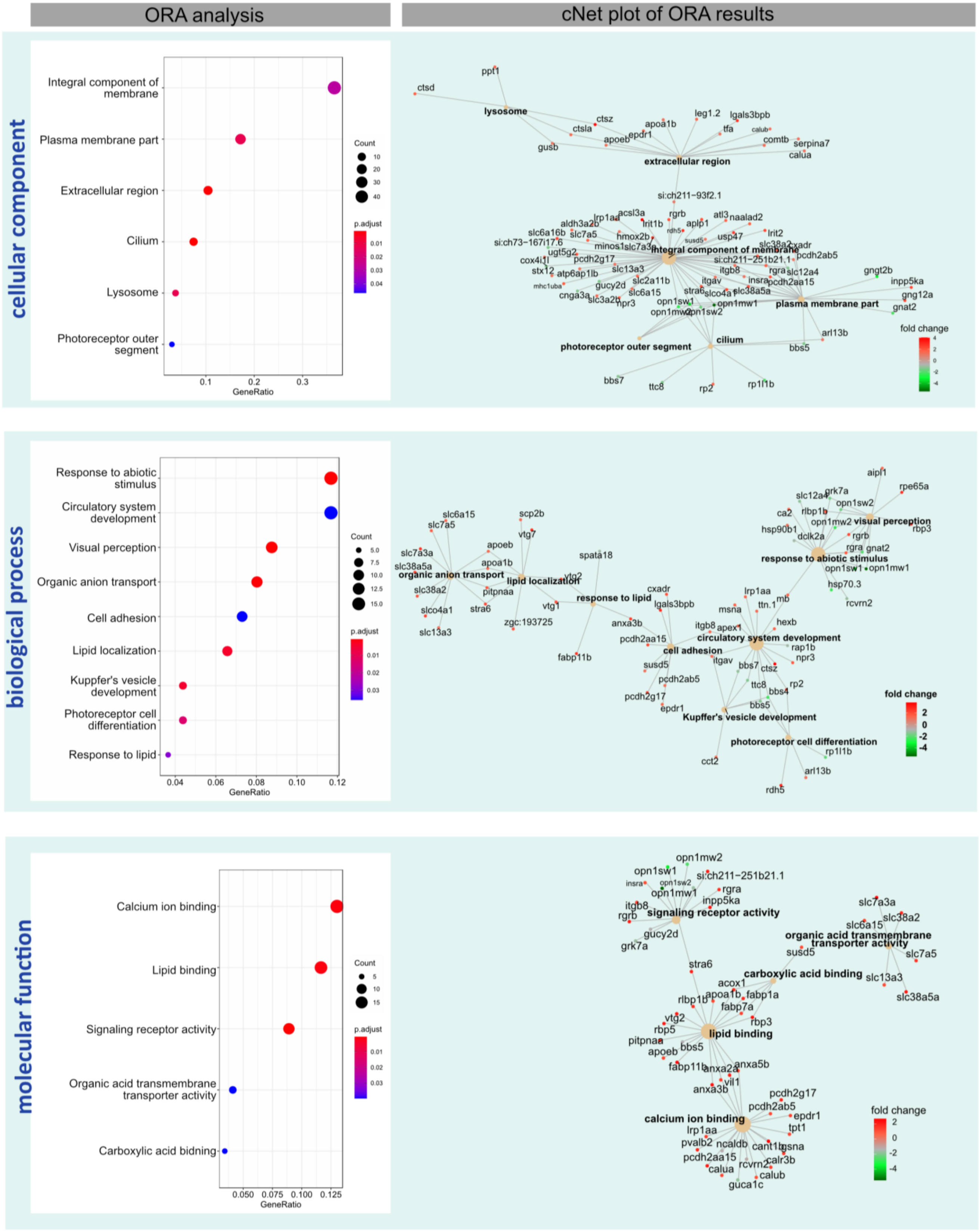

Over-representation of all significant GO-terms found in the cellular component (top), biological process (middle) and molecular function (bottom) using the proteomic dataset (adj-pval <0.05 & FC>±2). Terms that are significantly over-represented are shown in the dot plot (**left plots**). cNet plots (**right plots**) show the network of genes annotated to an over-represented GO-term. For details on the statistics, please see Supp. material and methods, data processing.

Suppl. Fig. 10: **Photoreceptors but not RPE cells express ApoE at 6 dpf**

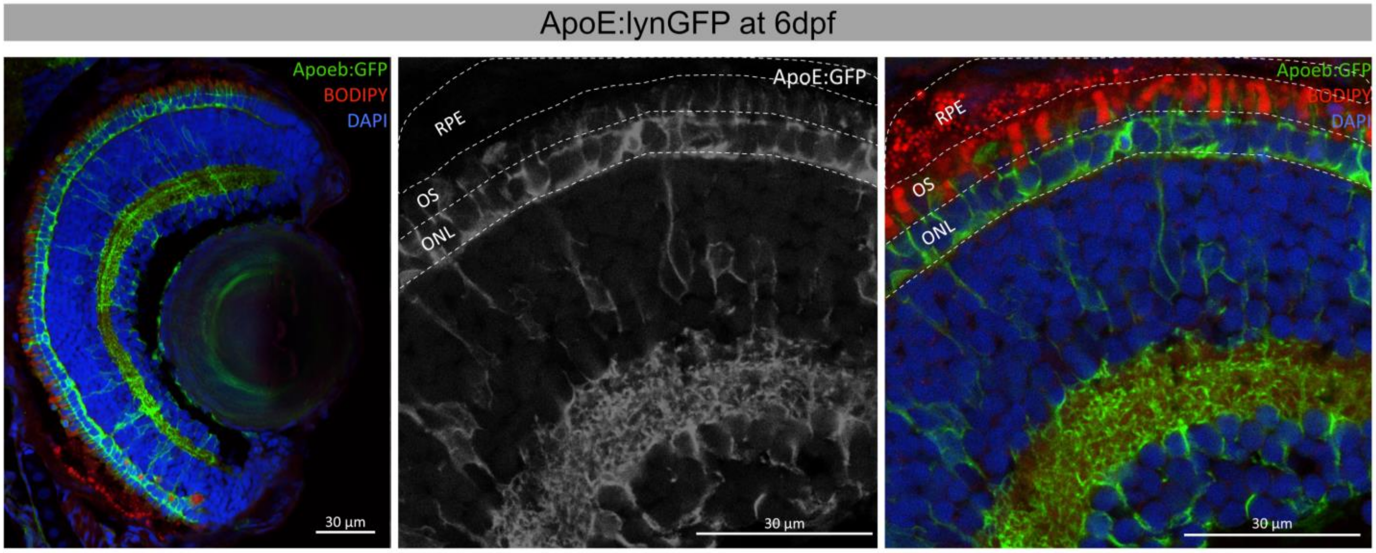


ApoE:lynGFP expression in a retinal section of a 6 dpf transgenic fish. GFP signal is found in photoreceptors and most likely Müller glia cells, but not in RPE cells. Scale bar: 30 µm; Abbreviations: *OS*, Outer segment; *RPE*, retinal pigment epithelium, *ONL* outer nuclear layer.

Suppl. Fig. 11: **Total serum cholesterol is unaffected in adult *zgbbs1* mutants**.

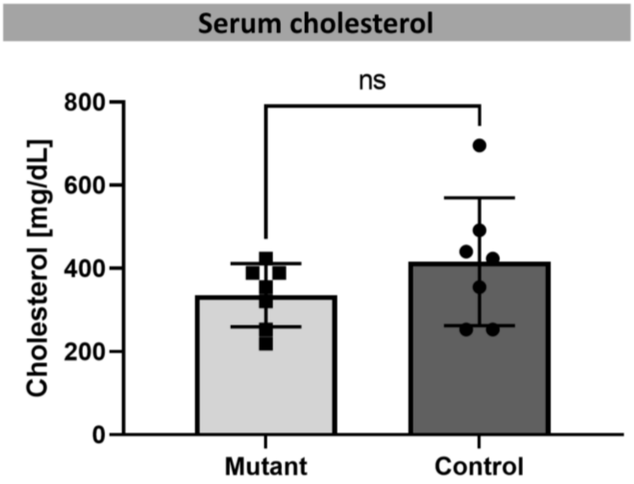


The blood serum of 5-month old *zgbbs1* mutant fish shows no increase in total cholesterol levels compared to age matched wild-type controls. This indicates that the cholesterol accumulation in the eye originates from a local distribution defect and not from systemic hypercholesterolemia. Statistics: independent two-tailed t-test, ns=p-value>0.05, Error bars show the standard deviation, sample size: (n=7 ctrl, n=7 Mut).

#### Supplementary tables

##### Suppl. Table 1: RNAseq_Dataset

Table of all genes found in the transcriptomic investigation with their normalized read counts and different expression analysis.

##### Suppl. Table 2: Proteomics_Dataset

Table of all detected proteins containing MAXquant intensities and different expression results

##### Suppl. Table 3: Dre_*bbs1-/-*_vs_*Bbs17-/-*_mouse_OS_Proteome

Table showing the direct comparison of the OS proteomic results from the zebrafish *bbs1* dataset presented in this work with the mouse Bbs17 mutant dataset published by Datta et al. 2015.

##### Suppl. Table 4: Lipidomic_Dataset

Table of all detected lipids of 5 month old mutant OSs. Raw data and processed data are shown in different worksheets.

##### Suppl. Table 5: Detailed statistics_ significance values

Table of the precise p-values, degrees of freedom, T-ratio and q-values of each multiple T-test.

#### Supplementary methods

**CRISPR/CAS9 Gene Editing:**

CRISPR guide RNAs were designed with ChopChop software. Bbs1 specific sgRNA was prepared as previously described[1]. In brief, we first performed a cloning-free PCR using a 60 bp gene specific oligo (TAATACGACTCACTATAGGACGTCTGGTGTCAGGCCAGTTTTAGAGCTAGAAATAGCAAG) containing one sp6 promoter for in vitro translation, 20 base spacer region specific to bbs1 target side, and one overlap region that anneals one constant oligonucleotide. Both *bbs1* specific and constant oligonucleotides (AAAAGCACCGACTCGGTGCCACTTTTTCAAGTTGATAACGGACTAGCCTTATTTTAACTTGCTATTTCTAGCTCTAAAAC) were annealed, fill-in with T4 DNA polymerase, cleaned with a PCR clean-up column, and transcribed into a sgRNA with the Ambion MEGAshortscript sp6 transcription kit.

For CRISPR/Cas9 injections 600ng Cas9 protein (GeneArt Platinum Cas9 Nuclease, Invitrogen) and 300 ng sgRNA were mixed in a total volume of 3 µl containing phenol red as injection control. Injections were performed as described before[2]. The web interface PCR-F-Seq q (http://iai-gec-server.iai.kit.edu) was used to assess the cutting efficiency of gRNA[3]. Bbs1 mutations were identified by Sanger sequencing and the founder fish was outcrossed to AB wild-type to balance potential off-target effects.

**Synteny and phylogenetic analysis:**

Synteny analysis of the zebrafish locus was done using the synteny database setting Homo Sapiens as outgroup (Variant: Ens70) (http://syntenydb.uoregon.edu/synteny_db/) as previously described[4]. For details please see supplementary methods. Parameters were adjusted to a sliding window size of 50 genes, and several genes in the vicinity of *bbs1* were used for additional syntenic comparison. Color and size of the final synteny graph was adjusted using affinity designer. The protein homology was calculated using Unipro UGENE (vers. 36.0) using standard methods from the Unipro UGENE Manual (vers. 36). In brief: Protein alignment was performed using T-coffee with the standard settings. The sequence homology was assessed based on amino acid similarity and the phylogenetic tree was build by the PHYLIP neighbour joining distance matrix using the Jones-Taylor-Thornton model.

**RNA sequencing:**

Library preparation was performed using the TruSeq Stranded Total RNA Library Prep Gold (Illumina, Inc, California, USA) after ribosomal RNA depletion. In brief, the quality of the isolated RNA was determined with a Fragment Analyzer (Agilent, Santa Clara, California, USA). Only those samples with a 260 nm/280 nm ratio between 1.8–2.1 and a 28S/18S ratio within 1.5–2 were further processed. The TruSeq Stranded Total RNA Library Prep Gold (Illumina, Inc, California, USA) was used in the succeeding steps. Briefly, total RNA samples (100-1000 ng) were depleted with ribosomal RNA and then reverse-transcribed into double-stranded cDNA. The cDNA samples were fragmented, end-repaired and adenylated before ligation of TruSeq adapters containing unique dual indices (UDI) for multiplexing. Fragments containing TruSeq adapters on both ends were selectively enriched with PCR. The quality and quantity of the enriched libraries were validated using the Fragment Analyzer (Agilent, Santa Clara, California, USA). The product is a smear with an average fragment size of approximately 260 bp. The libraries were normalized to 10nM in Tris-Cl 10 mM, pH8.5 with 0.1% Tween 20.

**Isolation and enrichment of photoreceptor outer segments**:

Photoreceptor outer segments of 5 month old zygotic adult fish were mechanically isolated using a modified version of the established techniques [5]. The OSs of 7 retinas were pooled for the proteomic analysis and of two retinas for the lipidomic analysis. Briefly, 5 month old dark adapted (>2h) zebrafish eyes were dissected under red light in the dark. The eye was cut open in 1x PBS (pH= 7.4) and the retina was slowly removed as little RPE as possible sticking to the retina. Removal of the RPE from the retina was done using a fine forceps. The retina was then transferred to a SYLGARD 184 silicone elastomer plate (Dow, Horgen, Switzerland) and flattened on the plate in a 30µL ice cold 1x PBS drop containing cOmplete, Mini, EDTA-free Protease-Inhibitors (Roche, Switzerland) OSs facing upwards. The outer segments were removed by gently swabbing over the OS using an extra fine, paintbrush. The rest of the retina was disposed and the 30µL PBS drop containing the OSs was transferred to a low-binding tube and kept on ice. The OSs of 7 retinas were pooled for the proteomic analysis and of two retinas for the lipidomic analysis. The OSs fraction was slowly pipetted onto a non-continuous sucrose gradient (47%, 37% and 32% Sucrose-PBS) and the tubes were washed 2x with 500µL ice cold 1x PBS. The tube was then carefully transferred to a pre-cooled 5810R centrifuge (Eppendorf AG, Hamburg Germany) with a free-swing bucket and centrifuged at maximum speed at 4°C for 120min, until the OS-fraction was clearly separated into the different fractions. The OS-containing fraction at the junction of 37% and 32% sucrose-PBS layer was removed and diluted in 1x PBS to a final concentration of sucrose-PBS <15%. The solution was spun down at 22'000rpm at 4°C in a 5424R centrifuge (Eppendorf AG, Hamburg Germany) and the supernatant removed. The protein pellet was then sent for proteomics or resuspended in 60µL water for the lipidomic analysis.

**Liquid chromatography-mass spectrometry analysis**:

Mass spectrometry analysis was performed on a Q Exactive HF-X mass spectrometer (Thermo Scientific) equipped with a Digital PicoView source (New Objective) and coupled to a M-Class UPLC (Waters). Solvent composition at the two channels was 0.1% formic acid for channel A and 0.1% formic acid, 99.9% acetonitrile for channel B. For each sample 3 µl of peptides were loaded on a commercial MZ Symmetry C18 Trap Column (100Å, 5 µm, 180 µm x 20 mm, Waters) followed by nanoEase MZ C18 HSS T3 Column (100Å, 1.8 µm, 75 µm x 250 mm, Waters). The peptides were eluted at a flow rate of 300 nL/min by a gradient from 8 to 27% B in 85 min, 35% B in 5 min and 80% B in 1 min. Samples were acquired in a randomized order. The mass spectrometer was operated in data-dependent mode (DDA), acquiring a full-scan MS spectra (350−1'400 m/z) at a resolution of 120'000 at 200 m/z after accumulation to a target value of 3'000'000, followed by HCD (higher-energy collision dissociation) fragmentation on the twenty most intense signals per cycle. HCD spectra were acquired at a resolution of 15'000 using a normalized collision energy of 25 and a maximum injection time of 22 ms. The automatic gain control (AGC) was set to 100'000 ions. Charge state screening was enabled. Singly, unassigned, and charge states higher than seven were rejected. Only precursors with intensity above 250'000 were selected for MS/MS. Precursor masses previously selected for MS/MS measurement were excluded from further selection for 30 s, and the exclusion window was set at 10 ppm. The samples were acquired using internal lock mass calibration on m/z 371.1012 and 445.1200.

The mass spectrometry proteomics data were handled using the local laboratory information management system (LIMS) [6] and all relevant data have been deposited to the ProteomeXchange

Consortium via the PRIDE (http://www.ebi.ac.uk/pride) partner repository with the data set identifier PXD026646.

**Data processing**:

The acquired raw MS data were processed by MaxQuant (version 1.6.2.3), followed by protein identification using the integrated Andromeda search engine. Spectra were searched against a Uniprot zebrafish reference proteome (taxonomy 7955, canonical version from 2019-07-01), concatenated to its reversed decoyed fasta database and common protein contaminants. Carbamidomethylation of cysteine was set as fixed, while methionine oxidation and N-terminal protein acetylation were set as variable modifications. Enzyme specificity was set to trypsin/P, allowing a minimal peptide length of 7 amino acids and a maximum of two missed cleavages. MaxQuant Orbitrap default search settings were used. The maximum false discovery rate (FDR) was set to 0.01 for peptides and 0.05 for proteins. Label-free quantification was enabled, and a 2-minute window for match between runs was applied. In the MaxQuant experimental design template, each file is kept separate in the experimental design to obtain individual quantitative values.

Protein fold changes were computed based on peptide intensity values reported in the MaxQuant generated peptides.txt file, using linear mixed-effects models. Pre-processing of the peptide intensities reported in the peptides.txt file was performed as follows: intensities equal zero are removed, non-zero intensities were log2 transformed and modified using robust z-score transformation to remove systematic differences between samples. For each protein, a mixed-effects model was fitted to the peptide intensities using the R-package lme4 [lme4]. We used the following model formula: transformedIntensity ~ Background_ * Knockout_ + (1 | peptide_Id), to model the factors Background and Knockout as well as their interactions, and modelling the peptide measurements as random effects. Fold changes and p-values were estimated based on this model using the R-packge lmerTest [lmerTest]. Next, p-values are adjusted using the Benjamini and Hochberg procedure to obtain the false discovery rates (FDR). In order to estimate fold-changes of proteins for which mixed-effects model could not be fitted because of an excess of missing measurements, the following procedure was applied: The mean intensity of a peptide over all samples in a condition was computed. For the proteins with no observation in one condition, we imputed the peptide intensities using the mean of the 10% smallest average peptide intensities determined in step one. Then the fold changes between conditions were estimated for each peptide, and the median of the peptide fold change estimates was used to provide a per protein fold change. No p-values were estimated in this case.

Gene ontology over representation analysis was performed on all the highly significant proteins (adj. P-value <0.05 & FC>±2), comparing them to all detected proteins to find terms that were overrepresented. ClusterProfiler (vers. 3.10.1) in R[7] was used for the analysis and the over represented terms (BH adj. p-value <0.05) were simplified and visualized using the enrichplot package (vers. 1.2.0).

**Lipidomics:**

Lipid extraction was performed as previously described [8] with some modifications. The MMC solvent (methanol: methyl tert-butyl ether: chloroform, 4:3:3, v:v:v) was supplemented with the SPLASH mix internal standard and additional internal standards: d7-sphinganine (SPH d18:0), d7-sphingosine (SPH d18:1), dihydroceramide (Cer d18:0/12:0), ceramide (Cer d18:1/12:0), deoxydihydroceramide (Cer m18:0 12:0) deoxyceramide (Cer m18:1 12:0) and glucosylceramides (GluCer d18:1/8:0 and GlcCer d18:1 18:0

(d5)) (Avanti Polar Lipids). Lipids were separated using a XSelect CSH C18 column (100 mm x 2.1 mm, 2.5 µm particle size, Waters Corp.) and an Exion UHPLC pump (Sciex Pte Ltd). Mobile phase A consisted of acetonitrile/water (60:40, v:v) with 10 mM ammonium formate and 0.1 % formic acid. Mobile phase B consisted of isopropanol/acetonitrile (90:10, v:v) with 10 mM ammonium formate and 0.1 % formic acid. Chromatography was conducted at 400 µl/min with constant column temperature at 50°C. The column was equilibrated with 40 % B, increased to 43 % B over 2 minutes, to 50 % B at 2.1 minutes, 54 %B at 12 minutes, 70 % at 12.1 minutes, 99 % B at 18 minutes and re-equilibrated with 40 % B for 2 minutes. Mass spectrometry analysis was carried out using a QTRAP 6500+ mass spectrometer in MRM acquisition mode (Sciex Pte Ltd). Data integration and analysis was performed using the Skyline software package and the MetaboAnalyst Suite [9,10]. MetaboAnalyst (vers. 5.0) was used to compare the median normalized lipid profiles of mutants and controls [11].
